# Spatio-temporal Proteomic Analysis of Stress Granule Disassembly Using APEX Reveals Regulation by SUMOylation and Links to ALS Pathogenesis

**DOI:** 10.1101/2020.01.29.830133

**Authors:** Hagai Marmor-Kollet, Aviad Siany, Nancy Kedersha, Naama Knafo, Natalia Rivkin, Yehuda M. Danino, Tsviya Olender, Nir Cohen, Thomas Moens, Adrian Higginbottom, John Cooper- Knock, Chen Eitan, Beata Toth Cohen, Ludo Van Den Bosch, Paul Anderson, Pavel Ivanov, Tamar Geiger, Eran Hornstein

## Abstract

Cytoplasmic stress granules (SG) form in response to a variety of cellular stresses by phase-separation of proteins associated with non-translating mRNAs. SG provide insight into the biology of neurodegeneration, including amyotrophic lateral sclerosis (ALS) because they approximate some of the molecular conditions for nucleation of insoluble aggregates in neuropathological inclusions. Whereas much has been learned about SG formation, a major gap remains in understanding the compositional changes SG undergo during normal disassembly and under disease conditions. Here, we address this gap by proteomic dissection of SG temporal disassembly sequence, using multi-bait APEX proximity-proteomics. We discover 109 novel SG-proteins and characterize at proteomic resolution two biophysically distinct SG substructures. We further demonstrate that dozens of additional proteins are recruited to SG specifically during disassembly, indicating that it is a highly regulated process. The involved proteins link SG disassembly, to mitochondrial biology and the cytoskeleton. Parallel analysis with C9ORF72-associated dipeptides, which are found in patients with ALS and frontotemporal dementia, demonstrated compositional changes in SG during the course of disassembly and focused our attention on the roles SUMOylation in SG disassembly. We demonstrate that broad SUMOylation of SG-proteins is required for SG disassembly and is impaired by C9ORF72-associated dipeptides, representing an unexplored potential molecular mechanism of neurodegeneration. Altogether, out study fundamentally increases the knowledge about SG composition in human cells by dissecting the SG spatio-temporal proteomic landscape, provides an in-depth resource for future work on SG function and reveals basic and disease-relevant mechanisms of SG disassembly.

**Highlights:**

- Multi bait APEX proximity labelling reveals 109 novel SG proteins and two distinct SG substructures.
- Proteomic dissection of SG temporal disassembly under basal conditions and with a model of neurodegeneration.
- Disassembly-engaged proteins (DEPs) include SUMO ligases that are recruited during normal disassembly and dysregulated in ALS-like conditions.
- Pervasive SG protein SUMOylation during SG disassembly is impaired by ALS-like conditions.

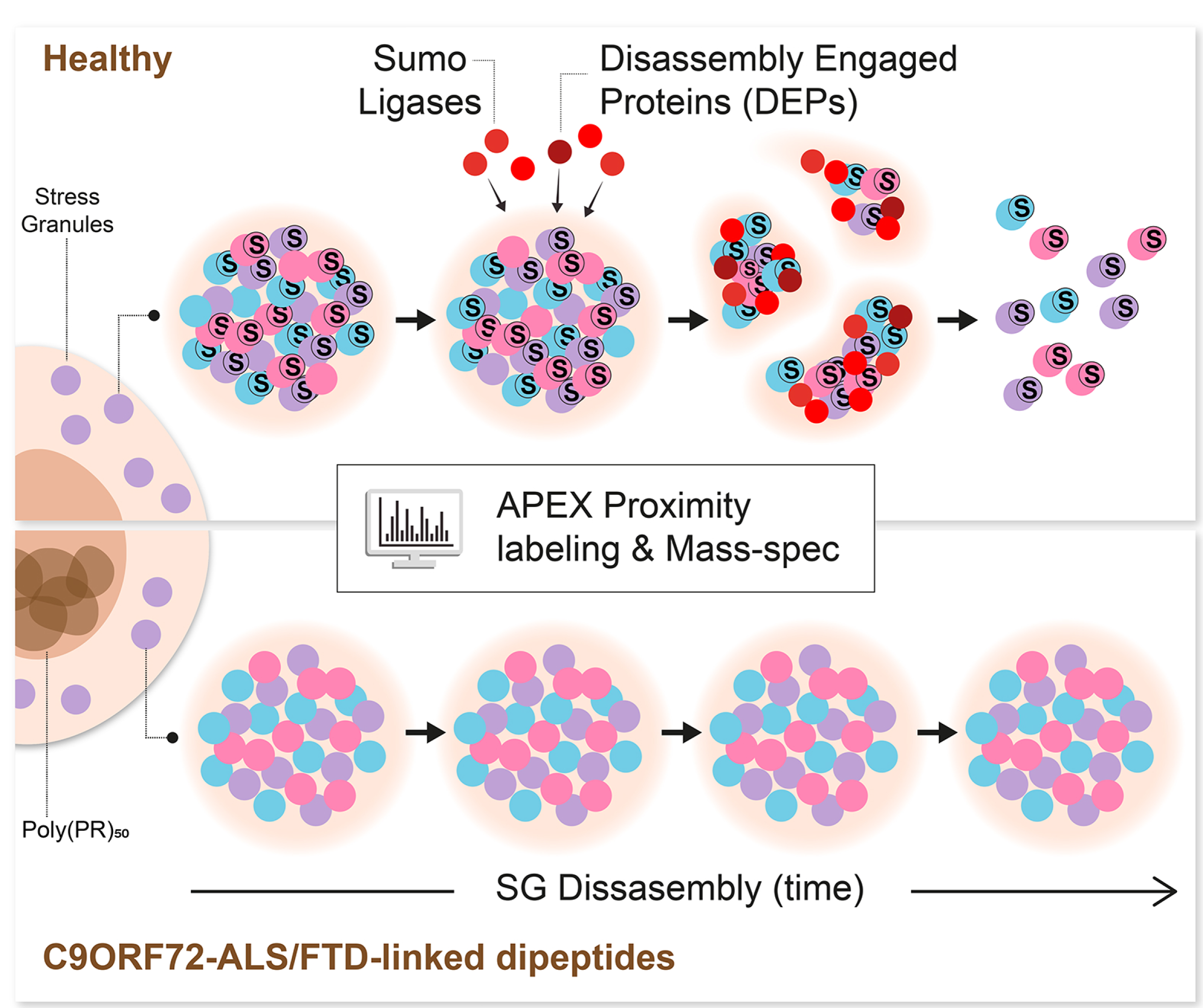

## Introduction

Stress granules (SG) are transient cytoplasmic membraneless condensates composed of ribonucleoproteins. They are formed through liquid-liquid phase separation of non-translating mRNAs and RNA-binding proteins via regulation of translation initiation machinery, in response to a variety of cellular stresses and dissolve upon return to normal growth conditions (Anderson and Kedersha, 2008, Decker and Parker, 2012, Buchan and Parker, 2009). The study of SG composition, as a means of gaining insight into their function, was first explored by biochemical fractionation and functional RNAi screens, which revealed hundreds of candidate SG proteins (Ohn et al., 2008, Jain et al., 2016). More recently, Proximity labelling proteomic analyses of SG uncovered cell-type and stress-specific SG composition (Markmiller et al., 2018, Youn et al., 2018).

Although the precise functions of SG remain to be determined, they are thought to play a protective role during cellular stress. However, persistent and abnormal SG may nucleate insoluble aggregates that are associated with human neurodegenerative diseases (Li et al., 2013), such as amyotrophic lateral sclerosis (ALS), a neurodegenerative disease of the human motor neuron system (Alberti and Dormann, 2019, Shorter, 2019, Taylor et al., 2016, Wolozin and Ivanov, 2019).

Thus, RNA-binding proteins that are residents of SG, such as TDP-43, HNRPNA2/B1 and FUS are encoded by genes that are mutated in different forms of ALS and are also found in neuropathological inclusions, in the brain and spinal cord of ALS patients (Molliex et al., 2015, Li et al., 2013, Ramaswami et al., 2013). In addition, the most common genetic form of ALS and of frontotemporal degeneration (FTD) is C9ORF72 disease is characterized by repeat associated non-ATG (RAN) translation of C9ORF72-ALS/FTD-linked dipeptides, that have been shown to affect SG formation (Boeynaems et al., 2017, Freibaum and Taylor, 2017, Zhang et al., 2018).

Intrinsically disordered regions (IDRs) underlie RNA-binding protein capacity to phase separate and condense SG. Accordingly, ALS-causing mutations are prevailing in IDRs and are thought to alter phase separation propensities and SG dynamics (Dormann et al., 2010, Kim et al., 2013, Molliex et al., 2015, Patel et al., 2015, Boeynaems et al., 2017, Lee et al., 2016, Hofweber et al., 2018, Elden et al., 2010, Wolozin and Ivanov, 2019). Post-translational modifications (PTMs), are preferentially affecting IDRs (Bah and Forman-Kay, 2016). Accordingly, PTMs, such as phosphorylation, Ubiquitination, poly-ADP ribosylation (PARP), and arginine methylation were shown to regulate SG assembly and phase separation (Kedersha et al., 1999, Leung et al., 2011, Babu et al., 2012, Han et al., 2012, Li et al., 2012, Banani et al., 2016, Dao et al., 2018, Sharkey et al., 2018, Hofweber et al., 2018). Moreover, SUMOylation of eIF4A2 was shown to contribute to SG formation (Jongjitwimol et al., 2016).

Whereas much has been learned about how SG form, many questions pertaining to basic SG biology, particularly the mechanisms underlying disassembly, and how they might be linked to disease, remain unanswered. Several pathways, including autophagy and ubiquitin-proteasome system were implicated in SG disassembly and ALS associated mutations were discovered in genes related to these mechanisms (Buchan et al., 2013, Seguin et al., 2014, Ganassi et al., 2016, Hjerpe et al., 2016, Wheeler et al., 2016, Protter and Parker, 2016, Turakhiya et al., 2018, Wang et al., 2019, Ivanov et al., 2019). However, a major gap in our understanding remains about the compositional changes SG undergo during the process of disassembly and its link to disease conditions.

APEX proximity proteomics has proven extremely useful in uncovering new facets of RNA granules (Lam et al., 2015, Lobingier et al., 2017, Markmiller et al., 2018, Liao et al., 2019). Here, we performed an APEX proximity proteomic study with three independent SG baits to characterize the constitutive SG proteome in unprecedented resolution: we identified over 109 novel SG proteins and the composition of substructures within SG. Furthermore, a temporal analysis of SG disassembly uncovered the existence of a group of proteins that are selectively recruited upon disassembly, and the impact of C9orf72 proline–arginine dipeptides that are associated with ALS and FTD (C9-ALS dipeptides). These data revealed a SUMO-dependent mechanism for the control of SG that is impaired by C9-ALS dipeptides, thus providing new insights into the patho-mechanisms of ALS.

## Results

### Comprehensive SG proteomic analysis by multi-bait APEX proximity labelling reveals 109 additional SG proteins

We characterized the protein composition of SG in living human cells, by means of APEX proximity labelling. APEX proximity labelling involves an engineered ascorbate peroxidase (APEX2), which is fused to a bait protein of interest. APEX2 facilitates free radical formation in its vicinity from H2O2, causing BP radical formation that tags biomolecules with a biotin moiety. Biotinylated proteins can be further purified over streptavidin beads and analysed by mass spectrometry (Hung et al., 2016). We hypothesized that using multiple APEX baits would allow a more comprehensive, and internally validated, characterization of SG beyond previous work (Markmiller et al., 2018, Jain et al., 2016, Youn et al., 2018). We engineered APEX fused to three RNA-binding proteins that have been extensively studied in SGs, namely, Ras GTPase-activating protein-binding protein 1 (G3BP1) (Aulas et al., 2015, Kedersha et al., 2016, Solomon et al., 2007, Tourriere et al., 2003, Cande et al., 2004, Markmiller et al., 2018), the Fragile X proteins, FMR1 (a.k.a. FMRP) and its autosomal homolog FXR1 (Zhang et al., 1995), downstream of a tetracycline-inducible promoter. We also fused a nuclear export signal (NES) to APEX to serve as a cytoplasm reference. To avoid overexpression artefacts including aberrations of organelle assembly dynamics or size (Anderson and Kedersha, 2008), tet-inducible FMR1-APEX and FXR1-APEX constructs were transfected to CRISPR-edited U2OS cells lacking FMR1/FXR1/FXR2 (Smith et al., 2019), whereas a tet-inducible G3BP1-APEX construct was transfected to a previously described U2OS line lacking G3BP1/G3BP2 (Kedersha et al., 2016). We selected single clones that displayed comparable construct expression and titrated induction by tetracycline to approximate endogenous expression levels and to be comparable across all baits (Fig. S1 A,B).

All three SG-APEX baits were activated by the introduction of the APEX substrates, biotin phenol (BP) and hydrogen peroxide (H2O2), which appropriately demarcated SG only when cultures were stressed with sodium arsenite (NaAsO2 300mM). Immunostaining revealed that the biotin signal co-localized with endogenous TIA1, an established SG marker (Fig. 1A) and NES localized correctly to the cytoplasm (Fig. S1 C). We confirmed that endogenous proteins were tagged with biotin by Western-blot analysis of pulled-down biotinylated proteins (Fig. S1 D,E). Importantly, biotinylation was specificlly observed only when the APEX substrates were introduced (Fig. S1 D,E,F).

**Figure 1.**
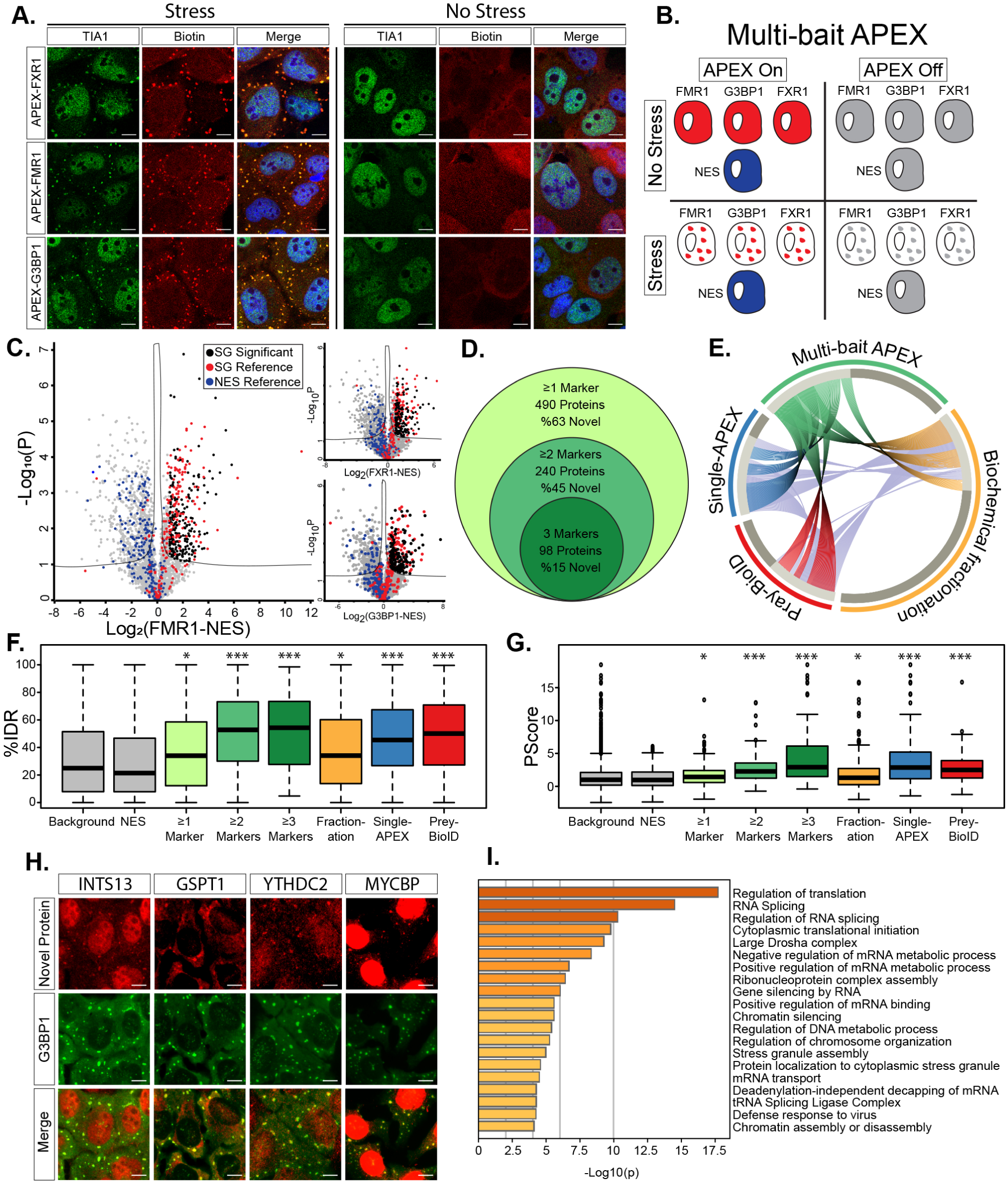
The proteome of Stress granules revealed by multi bait proximity labelling. (A) Confocal micrographs depicting FXR1-APEX, FMR1-APEX or G3BP1-APEX activity in U2OS cells, ± sodium arsenite stress (NaAsO2 400µM, 30 min.). Immuno-fluorescence depiction of TIA1, neutravidin-Texas-red staining of biotinylated proteins at the proximity of the APEX bait, and merged signal demonstrating the precise localization of the APEX activity in SGs. Lens x63, scale-bar 10µ. (B) Diagram of experimental design. Study with U2OS cells that stably express FXR1-APEX, FMR1-APEX, G3BP1-APEX or cytoplasmic NES-APEX. SG baits are diffusible in the cytoplasm without stress. NES-APEX remains diffusively cytoplasmic under stress conditions (NaAsO2 400µM, 30 min.), while FXR1-APEX, FMR1-APEX, G3BP1-APEX are recruited to SGs. APEX On: APEX peroxidase activity, induced by H2O2, causes BP radical formation that tags biomolecules in the bait vicinity with a biotin moiety. APEX Off: control for nonspecific activity without BP. Experiment performed in triplicates. (C) Volcano plot of relative protein levels in SG APEX relative to NES-APEX samples under stress conditions (x-axis log2 scale), analysed by MS. Y axis depicts the differential expression P values (−log10 scale). Black - novel SG proteins above specific marker cut-off (FMR1 = 0.96, FXR1 = 0.89, G3BP1 = 0.88) relative to NES. Student t-test with correction to multiple hypothesis by FDR. adj. P < 0.05. Red /blue - previously known SG proteins / cytoplasmic organellar proteins. (D) Venn diagram of multi-bait SG analysis and embedded results, revealing proteins identified by at least a single SG bait (associated with FXR1 and / or FMR1 and or G3BP1), at least two baits, or with all three baits together. (E) Circos plot of proteome depicted by at least two baits in our multi-bait APEX study (green), Single bait G3BP1-APEX ((Markmiller et al., 2018), Biochemical fractionation (Jain et al., 2016), or indirect (pray) analysis of data from BioID studies (Youn et al., 2018). Substantial overlap with the previously-known SG proteome is accompanied by the discovery of 109 novel and internally cross-validated proteins. (F) Box plot of intrinsically disordered region enrichment (%IDR, by IUPred, (Meszaros et al., 2018)) in the data of the current study and others (Markmiller et al., 2018, Jain et al., 2016, Youn et al., 2018). Background - all proteins identified in our MS analyses. Upper and lower quartiles, and extreme points. Wilcoxon signed-rank test p<0.005 (G) Box plot of SG proteome propensity to phase separate (Pscore (Vernon et al., 2018)) in the data of the current study and others as in (F). Upper and lower quartiles, and extreme points. ANOVA with Tukey post-hoc test P<0.05. (H) Confocal micrographs of immune-fluorescent detection of INTS13, GSPT1, YTHDC2, MYCBP in U2OS cells under stress conditions, and co-localization with the G3BP1 SG marker. scale-bar 10µm. (I) Bar graph depicting the significance of enrichment in the top 20 gene ontology (GO) terms for the SG proteome (-log p value), by metascape (Zhou et al., 2019).

We conducted three independent APEX experiments with the three SG baits activated in parallel, under basal and stress conditions, and controlled for both cytoplasm diffusive APEX signal (NES) and non-specific streptavidin bead binding (No BP; diagram in Fig. 1B). We identified 5987 proteins across all samples by label-free quantitative proteomics, using our analysis pipeline (Fig. S2A) with very high correlation across experimental replicates (Fig. S2B). We excluded proteins that were bound non-specifically to streptavidin beads (APEX off samples, 806 proteins, Fig S2C). To define candidate SG-interacting proteins, we applied student *t*-tests (FDR cut-off P < 0.05) to proteins associated with SG-APEX samples, relative to NES-APEX samples (Fig. 1C), while setting a stringent Log2 Fold change cut-off per each bait (see methods). Summary proteomic data related to this study is detailed in Table S1 and raw data in Table S2.

Well-characterized SG proteins were clearly enriched in our data (including, e.g., TIA1, UBAP2L, CAPRIN1, PABPC1, FUS and ATXN2). Overall, the mass spectrometry (MS) analysis identified 215 proteins associated with G3BP1-APEX (novel: 104 proteins, 48%, relative to the benchmark list (Youn et al., 2018)), 342 proteins associated with FMR1-APEX (novel: 196 proteins, 57%) and 260 proteins associated with FXR1-APEX (novel: 127 proteins, 49%).

In total, 490 proteins were identified with at least one of the APEX baits (novel: 311 proteins, 63%), of which 240 were detected with at least two baits (novel: 107 proteins, 45%) and 98 proteins were identified by all three SG-APEX baits (novel: 15 proteins, 15%) (Fig. 1D). Comparison of our internally cross-validated data (i.e., 240 proteins, identified by ≥2 baits) with other studies that investigated SG composition using other methodologies (Jain et al., 2016, Markmiller et al., 2018, Youn et al., 2018), demonstrated correct re-identification of ∼50% of the proteins in our data, and underscore the novelty of previously-uncharacterized and internally cross-validated 109 SG proteins (Fig.1E). Intriguingly, 72 of these unexplored proteins emerge from association with both FMR1 and FXR1 (66%), whereas only 36 (34%) of the new proteins were cross validated by G3BP1 and wither FMR1 or FXR1. These data underscore the value of using new baits (in this case, FMR1 and FXR1) that were not used in the past.

Using IUPred and Pscore (Meszaros et al., 2018, Vernon et al., 2018), we quantified the predicted enrichment of intrinsically disordered regions (IDRs) and propensity to phase separate in our data. SG proteins displayed values above the proteins enriched in the cytoplasm control (NES) or all proteins identified by the MS (Background); (P < 0.005 by Wilcoxon signed-rank test, P < 0.05 by ANOVA with Tukey post-hoc test, respectively, Fig. 1F, G). Moreover, IDRs and Pscore enrichment were significantly higher for proteins discovered by at least two or three baits, compared to the background, and resonated with IDR and Pscore values for protein lists from studies done by other methodologies. We were able to demonstrate the enrichment of 15 novel proteins in SG using immunofluorescence microscopy (Fig. 1H and Table. S5).

Gene ontology enrichment analysis (Zhou et al., 2019) showed that SG proteins in our data are associated with multiple aspects of RNA metabolism, in accordance with the known properties of SG 7 (Fig. 1I). However, we unexpectedly observed enrichment in proteins associated with DNA or chromatin. Our data also expanded the list of m6A binding proteins resident in SGs, in accordance with a critical role for RNA methylation in SG assembly and in phase separation (Anders et al., 2018, Ries et al., 2019), and the lists of post-translational modifiers associated with Poly-ADP ribosylation, ubiquitination or phosphorylation, which were all shown to regulate SG dynamics.

Altogether, our comprehensive analysis of the SG proteome, along with the newly discovered 109 SG proteins, increases knowledge about SG composition in human cells and serves as an in-depth resource for the SG proteomic landscape.

### Multi-bait APEX analysis allows the characterization of a pre-stress assemblies and distinct SG substructures

Macromolecular assemblies of SG proteins are known to exist under normal growth conditions, prior to induction of cellular stress (Markmiller et al., 2018, Youn et al., 2018). Therefore, we characterized the macromolecular assemblies within proximity of soluble FMR1, FXR1 or G3BP1 under basal conditions. Out of the 490 proteins that we characterized in SG upon stress (Table S1), 153 proteins were significantly enriched with APEX-labelled FMR1, FXR1 or G3BP1, relative to NES-APEX samples in basal conditions (Fig. 2A-B). Noteworthy, 113 proteins were associated with G3BP1 in soluble pre-stress complexes, whereas only 55 and 41 proteins were identified in the proximity of FMR1 or FXR1, respectively (Fig. 2B). 30 proteins were associated with G3BP1 and either FMR1 or FXR1 in pre-stress conditions, including several proteins that are essential for SG formation (dubbed, ‘SG seed’ Fig. 2B,C). Moreover, bioinformatic analysis of the SG seed proteins revealed a Pscore that is significantly higher than the mature SG proteome (P < 0.0005, ANOVA with Tukey post-hoc test, Fig 2D). Taken together, previously reported sub-microscopic pre-stress complex (Markmiller et al., 2018, Youn et al., 2018), exist mainly at the vicinity of G3BP1 and not FMR1 or FXR1. Within it, a SG seed exists that has exceptional phase separation potential and is suggested to initiate SG formation.

**Figure 2.**
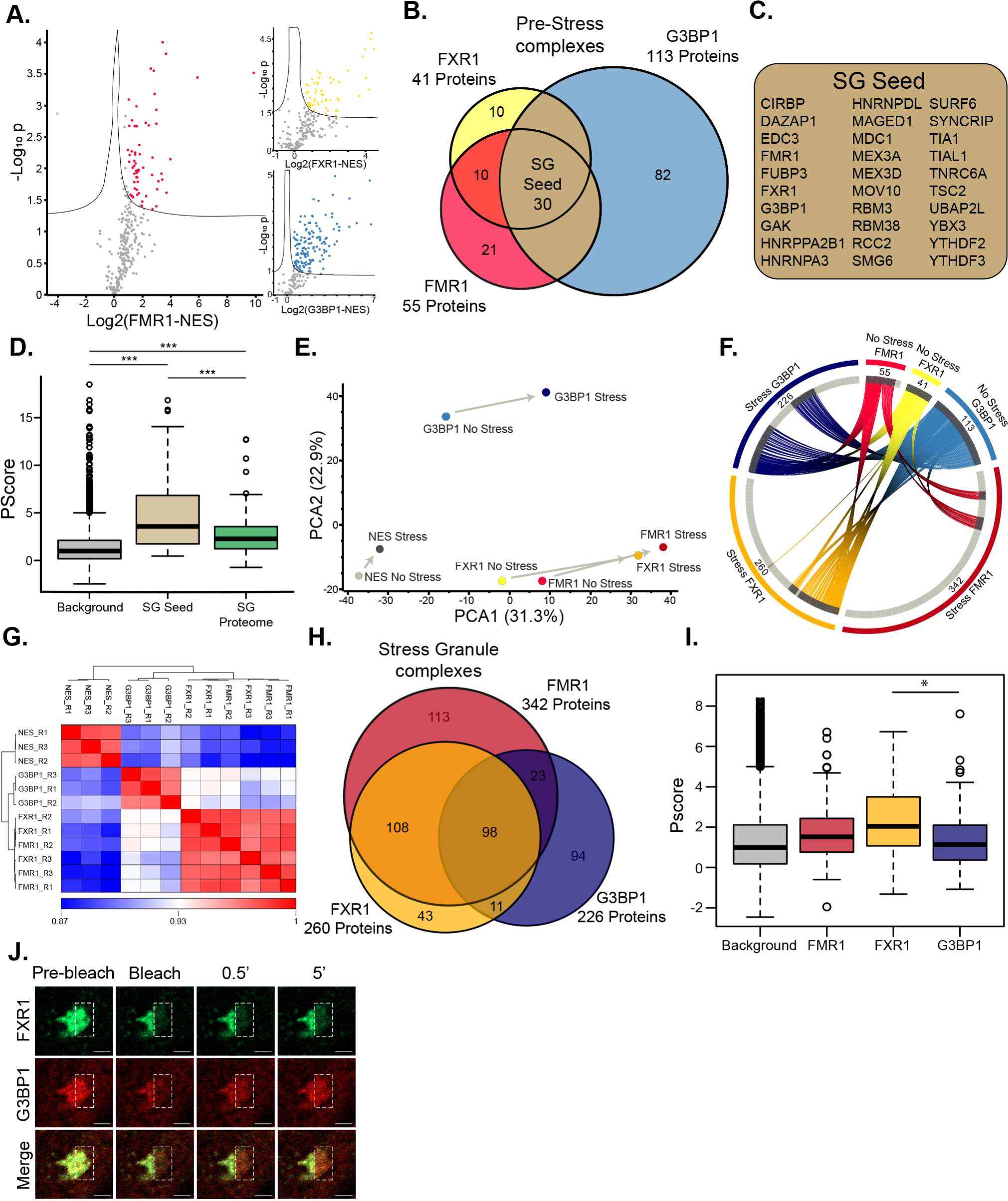
APEX study of proteome composition reveal the emergence of distinct SG substructures. (A) Volcano plot of relative protein levels in APEX samples relative to NES-APEX samples (x-axis log2 scale) under non-stress conditions. Y axis depicts the differential expression P values (−log10 scale). Black -proteins associated with APEX markers with at least 2-fold enrichment above values in NES. Student t-test with correction to multiple hypothesis by FDR. adj. P < 0.05. (B) Venn diagram of FMR1, FXR1 and G3BP1 proteomes under basal conditions, with number of proteins demarcated. (C) List of 30 SG seed proteins identified by 3 markers in basal, pre-stress, conditions. (D) Box plot analysis of propensity to phase separate (calculated by PScore (Vernon et al., 2018)) in the background (all proteins identified in our MS analyse, SG seed (30 proteins) and 240 proteins internally validated in mature SG. Upper and lower quartiles, and extreme points. ANOVA with Tukey post-hoc test P< 0.0005. (E) Principal component analysis (PCA) of FMR1, FXR1, G3BP1 and NES proteomes ± stress. Minimal compositional changes in the NES proteome influenced by stress (depicted as short PC vector), in contrast to more substantial compositional changes in the FMR1, FXR1, and G3BP1 proteomes. FMR1 and FXR1 proteomes collide. (F) Circos plot of proteins identified in basal and stress conditions, per each SG-APEX bait. ∼52% of the proteins associated with G3BP1 under stress, are already residents of the pre-stress G3BP1 complexes, whereas most of the proteins (84%) associated with FMR1 /FXR1 assemble *de novo* with stress. (G) Unsupervised clustering of Pearson correlation values for proteomes captured by SG APEX baits during stress. High similarity between FMR1 and FXR1 proteomes, which are distinct from the G3BP1 and NES proteomes. R1-3 experimental replicates. (H) Venn diagram of proteomes FMR1, FXR1, and G3BP1 proteomes in stress conditions, with number of proteins demarcated. (I) Box plot analysis of propensity to phase separate (Vernon et al., 2018) in the background (all proteins identified in our MS analyse, and proteins identified uniquely by only a single marker. ANOVA with Tukey post-hoc test P < 0.05. (J) FRAP analysis of U2OS cells that co-express G3BP1-RFP and FXR1-GFP. Laser bleaching was defined as time zero and snapshots were taken every 0.5 sec. The recovery of G3BP1-RFP was monitored at 0.5 min, whereas FXR1-GFP did not recover even after 5 min.

SG are non-homogenous and potentially contain substructures (Jain et al., 2016, Wheeler et al., 2016). However, little is known about the properties of such sub-regions. While FMR1, FXR1 and G3BP1 are all SG residents, principal component analysis of the proteomic data revealed that G3BP1 samples are distinct from FMR1/FXR1 samples under both basal and stress conditions (Fig. 2E). Notably, a substantial fraction of the proteins associated with G3BP1 under stress are already residents of the pre-stress G3BP1 complexes (∼52%), whereas most of the proteins associated with FMR1/FXR1 assemble only with stress (16%, 16% respectively), creating *de novo* a differentiated protein network that is distinct from that associated with G3BP1 (Fig. 2F).

In agreement with the PCA results, unsupervised clustering of the correlation between samples under stress revealed highly correlated mass spectrometric signatures associated with FMR1 and FXR1, which were dissimilar to the G3BP1 samples (Fig. 2G). Accordingly, the 108 proteins shared between FMR1 and FXR1 indicate a single substructure that is distinguishable from G3BP1 substructure (Fig 2H). Bioinformatic analysis revealed that proteins that were uniquely identified by FXR1 display significantly higher Pscore than proteins identified by G3BP1 (P < 0.05, ANOVA with Tukey post-hoc test, Fig 2I).

Because G3BP1 and FMR1/FXR1 may represent distinctive SG properties, we tested the hypothesis that they represent individual biophysical behaviours. We analysed SG liquidity by FRAP in cells that co-express mRFP-G3BP1 and GFP-FXR1. We observed rapid recovery of mRFP-G3BP1 after bleaching, whereas GFP-FXR1 displayed slow and incomplete recovery dynamics (Fig. 2J).

Together, proteomics and recovery kinetics reveal unexpected differences between G3BP1 and FMR1/FXR1 that may indicate the existence of distinct SG substructures with different biomaterial properties and proteomic composition. While one substructure organized around G3BP1 under basal conditions, and its composition modestly increases with stress, the other substructure assembles *de-novo* around FMR1/FXR1 in response to stress.

### Temporal analysis of SG disassembly reveals Disassembly Engaged Proteins

The compositional changes that SG undergo during the process of disassembly are poorly understood. To better understand normal SG disassembly, we first monitored disassembly by mRFP-G3BP1 live imaging (in the presence of GFP expression vector) after washing out medium with sodium arsenite (300uM, 30min). We noticed that microscopically-visible disassembly started at ∼40 min. after stressor washout and was completed in ∼120 minutes (Fig. 3 A). In this context, we then used APEX and MS to characterize the SG disassembly proteome. We chose FMR1-APEX as it exhibited the largest proximity proteome of the three SG baits and because, as opposed to G3BP1, it does not preferably bind to the ubiquitin-proteasome (UPS) system (Kedersha et al., 2016). From the MS results, we excluded proteins that were non-specifically bound to streptavidin beads, and normalized the protein intensity values of FMR1-APEX samples to the values measured in the NES-APEX. Out of the 7003 proteins that were identified in all MS samples, 584 were reproducibly enriched in the FMR1 samples, relative to levels in the NES samples by at least two-fold, in at least one of the time points (Fig. S3A pipeline description, ANOVA statistical test, FDR <0.05). Unexpectedly, 349 of the 584 proteins, were not detected in SG under sodium-arsenite stress and ascended to associate with SG only during the disassembly process (Table S3). We named these newly-recruited proteins, Disassembly Engaged Proteins (DEPs, Fig. 3B). The recruited DEPs suggest that SG disassembly is a highly regulated process. DEPs are associated with processes that were previously linked to the turnover of SG, including autophagy and ubiquitin pathways (examples in Fig. 3C and fully described in Table S3). In addition, heat-shock proteins, RNA helicases, cytoskeletal proteins, and mitochondrial proteins, were also observed. SG proteins were enriched with IDRs and displayed higher likelihood to phase separate than DEPs (P < 0.0005 by student *t*-test, P < 0.0005 by Wilcoxon signed-rank test, respectively, Fig. 3D,E). These data indicate that DEPs are not necessarily engaged to SG by phase separation and suggest that their programmed engagement may be governed by another mechanism that is currently unexplored.

**Figure 3.**
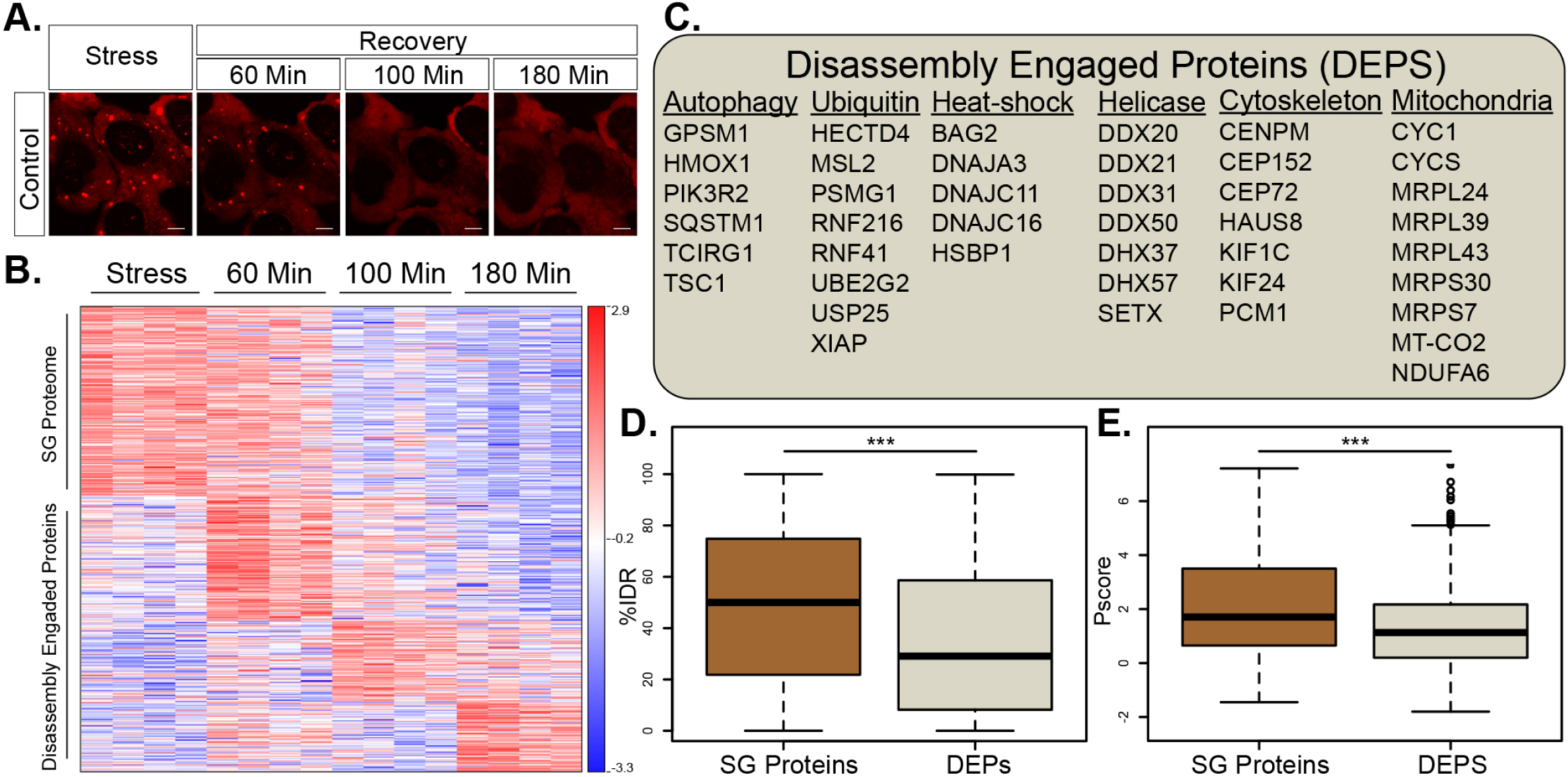
Temporal resolution of SG disassembly under normal conditions reveals a network of Disassembly Engaged Proteins. (A) Representative micrographs depicting RFP-G3BP1 in stressed U2OS cells and 60, 105 and 180 minutes during recovery, after the stressor was washed-out. (B) Heatmap of unsupervised clustering of proteins associated with SG disassembly. Proteins specifically enriched in SG relative to the cytoplasm, if exceeding a 2-fold enrichment in FMR1-APEX / NES-APEX and P < 0.05 by student t-test with correction to multiple hypothesis by FDR). 235 proteins are enriched in SG, while 349 proteins are recruited to SG only once stress is removed and disassembly dynamics ensue. (C) A list of representative Disassembly Engaged Proteins (DEPS) associated with different cellular pathways. (D) Box plot analysis of intrinsically disordered region enrichment (%IDR, (Meszaros et al., 2018)) in 235 SG resident proteins or 349 DEPs. Upper and lower quartiles, and extreme points. Wilcoxon signed-rank test P<0.0005. (E) Propensity to phase separate (Vernon et al., 2018), in 235 SG resident proteins or 349 DEPs. Upper and lower quartiles, and extreme points. 2-sided student *t*-test P< 0.0005.

### C9ORF72-ALS dipeptides alter the SG composition and recruitment of disassembly engaged proteins

It was suggested that SG serve as the origin of insoluble cytoplasmic aggregates in ALS (Li et al., 2013, Buchan et al., 2013). Thus, better understanding of SG disassembly dysregulation may reveal the mechanisms that nucleate inclusions in the diseased brain. Therefore, we stably expressed a tetracycline-inducible GFP protein fused to 50 proline-arginine di-peptide repeats (GFP–poly(PR)50 ; (Wen et al., 2014)), associated with the C9ORF72 subtype of ALS (Freibaum and Taylor, 2017, Tran et al., 2015, Wen et al., 2014). Expression of GFP–poly(PR)50 in U2OS cells resulted in the formation of nuclear aggregates, as previously reported (Freibaum and Taylor, 2017, Tran et al., 2015, Wen et al., 2014). These aggregates neither co-localized with SG, nor caused SG formation. However, a loss of ∼30% of the cultured cells was measured 3 days after induction of GFP–poly(PR)50 expression (Fig. S3B). GFP–poly(PR)50 expression resulted in aberrant SG disassembly, wherein 30% of SGs failed to dissolve into the cytoplasm even after six hours (ANOVA repeated measurement P < 0.0005, Fig. 4A,B), in accordance with the behaviour of other C9orf72 associated dipeptides (Zhang et al., 2018).

**Figure 4.**
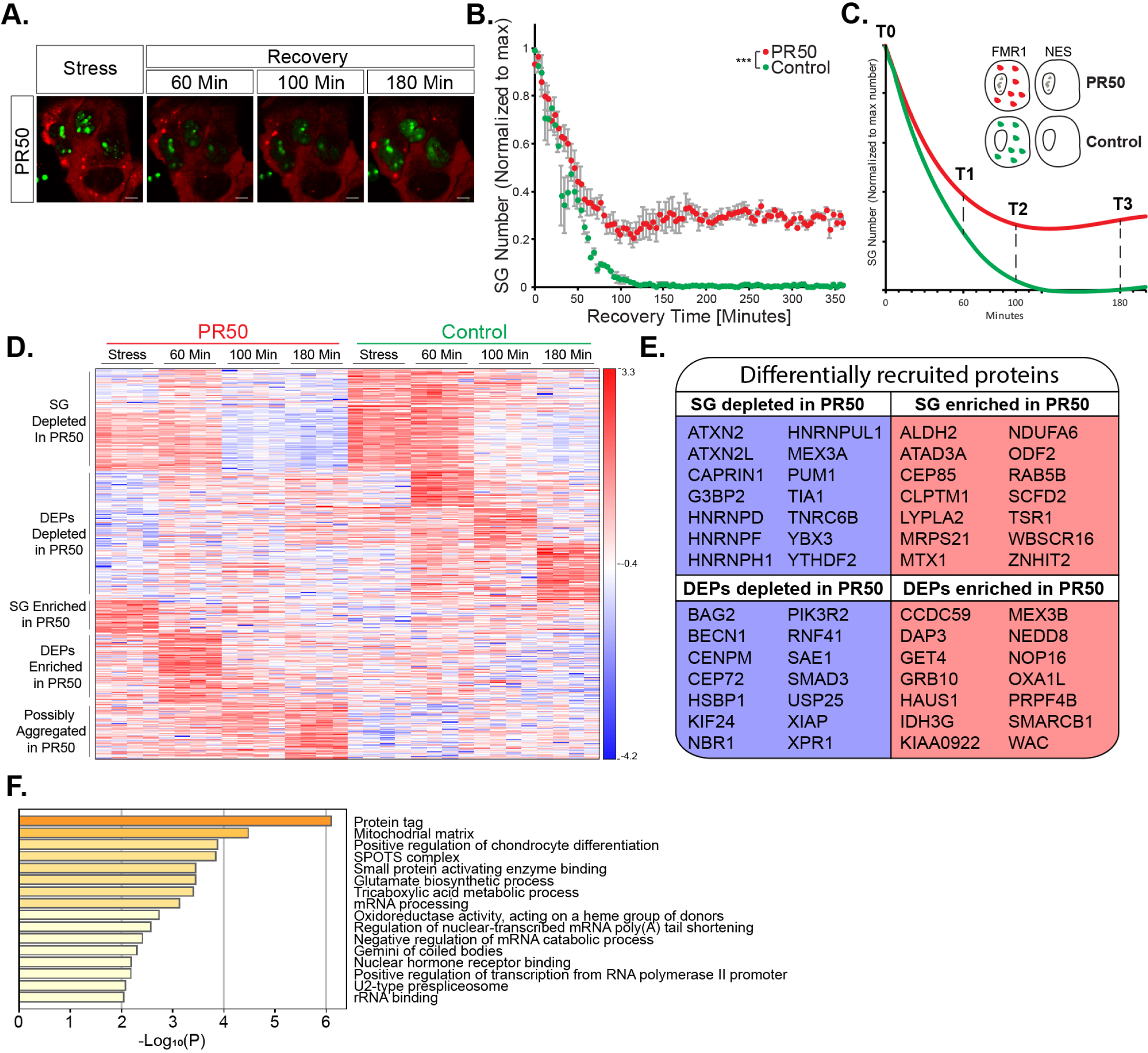
Temporal resolution of SG disassembly under normal conditions and with C9-ALS associated dipeptides. (A) Representative micrographs depicting RFP-G3BP1 in stressed U2OS cells that express GFP-poly(PR)50 and during recovery, after the stressor was washed-out. (B) Graph quantification of SG disassembly dynamics by live Cherry-G3BP1 imaging after stressor washout with inducible GFP (control) or GFP-poly(PR)50 expression (PR50). SG number, normalized to maximal SG numbers per field (y-axis) as a function of time after stress washout (x-axis). Three experimental repeats for measurement with 4 different areas / well. Representative experiment from > 3 independent live imaging studies. ANOVA repeated measurement P < 0.0005. (C) Diagram of study design. Inducible FMR1-APEX or NES-APEX baits, with inducible GFP (control) or GFP-poly(PR)50 expression (PR50). APEX proximity labelling activity was induced at T0 (stress) or at three time points after washout. (D) Heatmap of unsupervised clustering of proteins that were differentially associated with SG under normal and expression of GFP–poly(PR)50 during the course of disassembly. (E) Proteins specifically enriched in SG relative to the cytoplasm (FMR1-APEX / NES-APEX 2-fold change) and GFP–poly(PR)50 / GFP by 2 way ANOVA with FDR P < 0.05). 425 proteins that were differentially associate (176 enriched / 249 depleted) d with normal vs. GFP–poly(PR)50 SG in at least one of the time points in the series. (F) A table with a representative list of differentially recruited proteins. (G) GO term analysis of proteins depleted during SG disassembly (time 60min, 100 min, 180min), under GFP–poly(PR)50 conditions, relative to controls. Bar graph of -log10 of P-value, after 2-way ANOVA statistics.

In parallel with the proteomic analysis of normal disassembly sequences, that was described in Fig 3 (GFP control vector), we also performed APEX and mass spectrometry in the presence of GFP– poly(PR)50 (study design diagram Fig. 4C and pipeline analysis in Fig. S3A). Overall, we found 425 proteins that were differentially associated with normal vs. GFP–poly(PR)50 SG in at least one of the time points in the series. 176 / 249 were relatively enriched / depleted in GFP–poly(PR)50 conditions, respectively (Fig. 4D).

During stress, 59 proteins were relatively depleted from mature SG under GFP–poly(PR)50 conditions, including classical SG proteins (CAPRIN1, G3BP2, YBX3 and TIA1), while 32 proteins were relatively enriched (2 way ANOVA with contrast analysis and FDR correction P<0.05). When the stress was washed away to trigger SG disassembly, DEPs recruitment, was also inhibited by GFP–poly(PR)50 expression (Fig. 4D,E). Finally, gene ontology term analysis (Zhou et al., 2019) of proteins that were differentially inhibited by GFP–poly(PR)50 specifically during disassembly revealed the potential relevance of “protein tags” (Fig. 4F), suggesting the involvement of post-translational modifications in the regulation of SG disassembly.

### Pervasive SUMOylation of SG proteins controls SG disassembly, and is impaired by C9ORF72-ALS proline–arginine dipeptides

The ubiquitin-like SUMO ligase complex consist of E1, E2 and E3 ligases, which SUMOylate target proteins (Geiss-Friedlander and Melchior, 2007). We identified SAE1 (E1), UBE2I (E2), and the E3s, RANBP2 and TOPORS, as SG DEPs. Furthermore, SAE1 and UBE2I were significantly depleted from SG upon GFP–poly(PR)50 expression (Fig. 5A). We therefore hypothesized that SUMOylation is important for SG disassembly.

**Figure 5.**
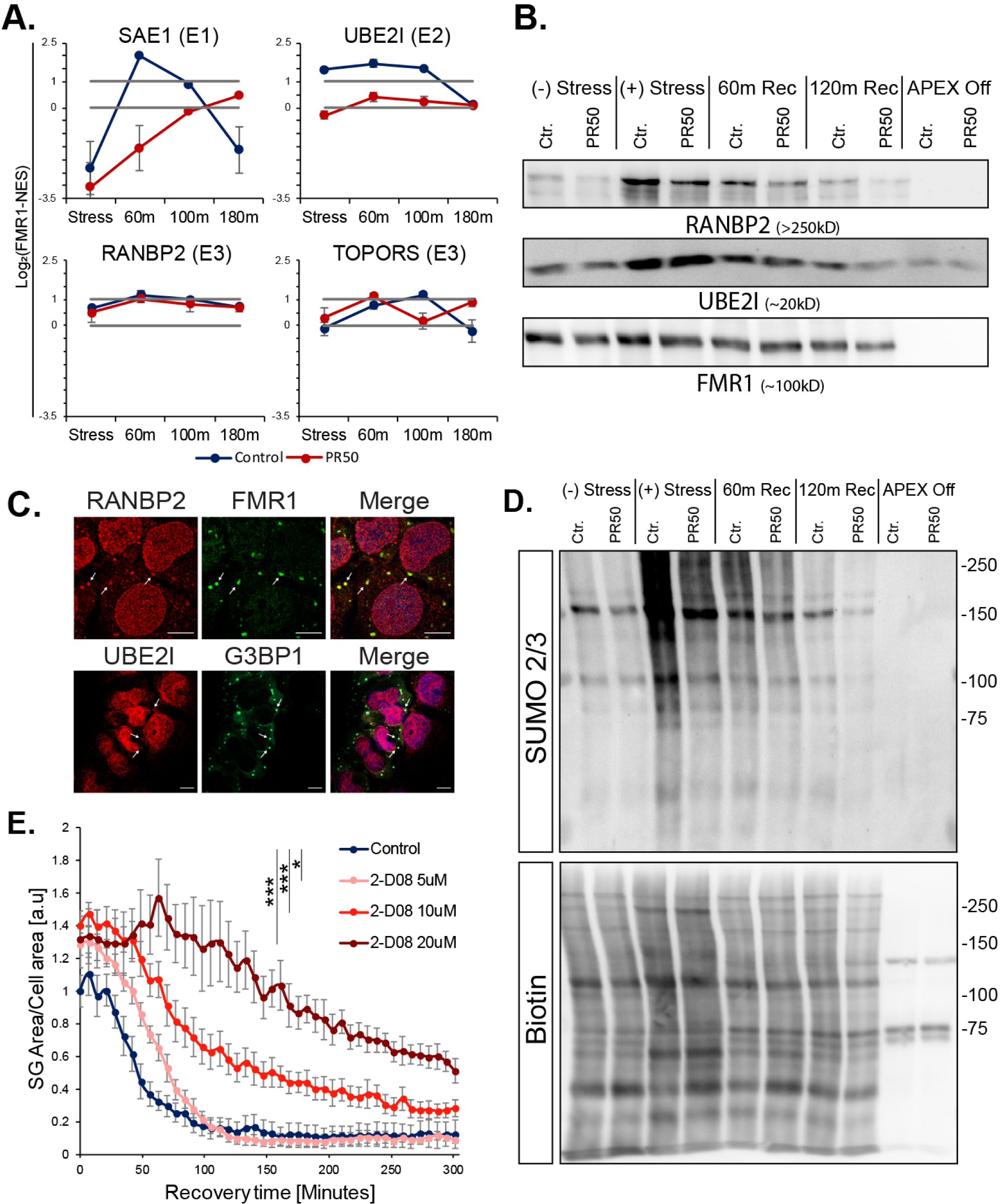
Pervasive SG SUMOylation controls SG disassembly. (A) Graph of MS quantification of UBE2I, SAE1, TOPORS, and RANBP2 in U2OS SGs. Log2 fold change of LFQ intensity in FMR1 minus NES, in stress conditions, and three time points after washout. Red-GFP–poly(PR)50. Lower bar - levels in the cytoplasm, higher bar-2-fold enrichment. Mean ± SEM. (B) Western blot analysis after FMR1-APEX activity and streptavidin pulldown of biotinylated SG proteins for detection of RANBP2, UBE2I, and FMR1-APEX as loading reference. RANBP2 and UBE2I are present in SG in response to stress and during recovery. GFP– poly(PR)50 conditions inhibit RANBP2 recruitment. (C) Immunofluorescence analysis of RANBP2 and UBE2I localization in SG. FMR1 or G3BP1 as SG markers. Merge includes demarcated nucleus (blue, DAPI). Lens x63. Scale bar - 10µm. (D) Western blot analysis after FMR1-APEX activation and streptavidin pulldown of biotinylated SG proteins for detection of SUMO 2/3 -conjugated proteins (upper blot) and loading control developed with streptavidin for detection of biotinylated proteins. Extensive SUMOylation of SG proteins seen as smear at 100-250kD and gradual decrease associated with disassembly. GFP–poly(PR)50 expression inhibit SUMOylation. Representative blot from >3 studies. (E) Graph quantification of SG disassembly dynamics by live GFP-G3BP1 imaging with increasing concentrations of 2D08, a SUMOylation inhibitor. Stress was induced with sodium arsenite (300 um, for 30 min), washed out, and 2D08 was introduced. SG area, normalized to cellular area (y-axis) as a function of time after stress washout (x-axis). Repeated measures ANOVA, P < 0.05. Three experimental repeats for measurement with 4 different areas / well. Representative experiment from > 3 independent live imaging studies.

First, we validated that UBE2I and RANBP2 are genuinely associated with SG in a stress-dependent manner by performing Western blot analysis on SG proteins that were pulled-down by FMR1-APEX (Fig. 5B). An immunofluorescence study further supported the presence of UBE2I and RANBP2 in SG (Fig. 5C). In addition, SG recruitment of RANBP2 was impaired by GFP–poly(PR)50 expression.

We then tested whether the SG proteome is SUMOylated in response to stress by performing FMR1-APEX pull-down and Western blot analysis of SUMO modifications. We observed a stress-dependent smear of SUMO1/2/3, primarily at protein sizes of 100-250kD, with reference to biotinylated proteins in the loading control blot (Fig. 5D and S4A). This study reveals broad SUMOylation of SG proteins, in a stress-dependent manner, and not exclusively limited to the process of disassembly. Additionally, conjugation of SUMO 2/3, but not of SUMO1 to SG, was impaired by GFP–poly(PR)50 expression (Fig. 5D).

To test whether SUMOylation plays a functional role in SG disassembly, we performed live SG imaging in the presence of the small molecule SUMO inhibitor, 2D08, which blocks the transfer of SUMO from the E2 SUMO ligase, UBE2I, to its substrates (Kim et al., 2014). 2D08 reduced overall SUMOylation, as is expected (Fig. S4B) and neither drove SG formation, nor the phosphorylation of eIF2 alpha (Fig. S4C,D). When 2D08 (at 5-20 µM) was administered for 30 minutes along with sodium-arsenite and included in the medium also after the stressor was washed-out, disassembly was attenuated in a concentration-dependent manner (ANOVA repeated measurement P < 0.0005, Fig 5E). Therefore, inhibition of SUMOylation is important for the disassembly process and is further reminiscent of the impact of GFP–poly(PR)50 expression.

## Discussion

Proximity proteomic mapping by multiple baits set new standards for membraneless organelles. Accordingly, our study can be used as in-depth database for the spatio-temporal landscape of SG. Particularly, we provide for the first time an analysis of SG disassembly kinetics at proteome resolution, revealing basic and disease-relevant mechanisms.

In agreement with previous studies (Markmiller et al., 2018, Youn et al., 2018), we observed sub-microscopic pre-stress complexes. However, we were able to additionally characterize these as likely only a single pre-stress entity in the proximity of G3BP1. More stringent analysis, by all three baits, revealed the identity of 30 proteins at the SG seed. Many of these are classic SG proteins collectively exhibit superior capacity to phase separate. The degree of similarity between SG seed and the SG Core that is isolated from mature SG (Jain et al., 2016, Wheeler et al., 2016) is currently a mystery, but considering that several SG seed proteins are necessary for SG formation and the high propensity to phase separate, we suggest that SG seed proteins may initiate SG assembly and perhaps develop into the Core of the mature granule.

In a similar context, (Wheeler et al., 2016) demonstrated substructures inside SG, but their proteome composition remains unexplored. The unique power of multi-bait APEX allowed us to characterize the proteome composition of two distinguishable substructures inside mature SG and FRAP analysis suggested that the two substructures display different biomaterial properties. Our interpretation is that one substructure is probably created by maturation of the SG seed that pre-exists under basal conditions at the proximity of G3BP1, and only modestly changes with stress. This perhaps is the SG Core (Wheeler et al., 2016). However, the other substructure assembles *de-novo* around FMR1/FXR1 in response to stress.

The reasoning that better understanding of SG disassembly could lead to the discovery of new pathways and therapeutic targets in neurodegeneration (Li et al., 2013), drove us to use APEX and mass spectrometry to temporally study the process of SG disassembly in normal and C9ORF7-ALS-like conditions, which impairs SG disassembly.

This analysis, which was impossible in the past, revealed a group of proteins that are recruited to SG only during disassembly, which we named disassembly-engaged proteins (DEPs). The existence of DEPs indicates that SG disassembly is a controlled process that occurs in a stepwise manner and cannot be described as passive dissolution of SG components into the cytoplasm. A substantial number of these DEPs are associated with processes that were previously linked to the turnover of SG, including autophagy and ubiquitin. Rather unexpectedly, mitochondrial proteins make the largest functional group of proteins that engage with SG during disassembly. The potential role of mitochondrial proteins in SG disassembly is unexplored and may suggest previously unappreciated links, tying cellular metabolism to SG disassembly or new moonlighting functions for mitochondrial proteins.

Many DEPs were aberrantly-recruited to SG when GFP–poly(PR)50 is expressed. Therefore, DEPs characterized in our studies are primary candidates for follow-up research on pathways that are potentially relevant to the pathogenesis of ALS and FTD. These include for example, BECN1, NBR1 and PIK3R2 that might highlight a particular involvement of mediators of autophagy.

In addition, several SUMO ligases are DEPs and hence connect SUMOylation to SG disassembly. Our study accordingly demonstrated broad, stress-dependent SUMOylation of SG proteins, and that SUMOylation activity is functionally important for SG disassembly. One compelling hypothesis about the SUMO-dependent SG disassembly may involve SUMO-targeted ubiquitin ligase (STUbL) family, which ubiquitinate direct the degradation of SUMO modified proteins (Sriramachandran and Dohmen, 2014).

The broad SUMOylation of SG proteins we observe during stress is consistent with the reported SUMOylation of eIF4A2 (Jongjitwimol et al., 2016) and suggest that many other proteins are SUMOylated. However, the dynamic application of a small molecule SUMO inhibitor, 2D08, suggest that *de-novo* SUMOylation is independently required for the process of SG disassembly.

GFP–poly(PR)50 expression impaired SUMO ligase recruitment and SG SUMOylation, suggesting that SUMO plays a part in the mechanism by which C9orf72 dipeptides attenuate SG disassembly. This can further represent an unexplored molecular mechanism of neurodegeneration acting via control of SG turnover.

Although our data are extensive and internally cross-validated, we were limited to simple cultured cells because proteomic analyses requires high yields and can thus not yet be performed, for example, in human motor neurons differentiated from iPSCs. When yield limitations are overcome by mass iPSC production, performing proteomics analysis in human motor neurons will become feasible and enable additional breakthroughs in the research of neurodegenerative diseases. The focus on SUMOylation as a follow up to our proteomic analysis, further encourages molecular studies to uncover the involvement of additional proteins suggested by our data.

Altogether, our study provides an in-depth resource for follow-up studies, dissecting the SG spatio-temporal proteomic landscape, and revealing basic and disease-relevant mechanisms of SG disassembly.

## Acknowledgements

EH is the Mondry Family Professorial Chair and Head of the Nella and Leon Benoziyo Center for Neurological Diseases. We thank Alice Ting (Stanford University) for creating and generously sharing the APEX proximity labelling technology, Davide Trotti (Jefferson University) for the 50x proline-arginine repeat vector whose insert serves as basis of the GFP–poly(PR)50 construct. We thank Dr. Vivek Advani (Harvard Medical School) for assistance with preliminary experiments. WE Thanks Assaf Kacen, Daoud Sheban, Avital Eisenberg Yifat Merbl (Weizmann Institute of Science) for advice and preliminary experiments. We thank Hornstein, Anderson and Ivanov lab members for helpful critiques and Life Science Editors for editorial assistance.

## Author contribution statement

HMK and EH conceived research and analysed the data. NK, PI, PA, provided valuable reagents for research. HMK, AS, NK, NR, YDM, BTC Performed tissue culture APEX proximity labelling molecular studies and microscopy. HMK, NK, TG Performed mass spectrometry. HMK, TO, NC, Performed computational data analysis. TM, LVDB, CE, JCK, AH, Assisted research. HMK and EH wrote the manuscript, with comments and final approval by all other authors. TG is corresponding author for MS, EH is the corresponding for all other facets of the work.

## STAR Methods

### Mammalian Cell Culture

Human Bone Osteosarcoma Epithelial Cells, U2OS (U-2 OS, ATCC HTB-96), were cultured in growth media consisting of Dulbecco’s Modified Eagle Medium (DMEM, Biological Industries, 01-050-1A) supplemented with 10% fetal bovine serum (FBS, Biological Industries, 04-001-1A), 1% penicillin-streptomycin (Biological Industries, 03-0311B) at 37°C, with 5% CO2. Stress granules induced by NaAsO2 (200-300 μM, Sigma-Aldrich, 71287).

### Cloning of APEX vectors

V5 epitope tag, SG bait protein (FMR1, FXR1 or G3BP1) and APEX2 were subcloned into pcDNA4.0-TetO vector downstream of Tet-On inducible promoter and transfected by using Lipofectamine 2000 Transfection Reagent (Thermo Fisher Scientific, Cat# 11668027) to U2OS cells that express the Tet-Repressor protein. Antibiotic resistance selection against Zeocin (250µg/ml, Invivogen ZEL-41-01) or puromycin (2ug, Invivogen, ANT-PR)) enabled the isolation of single cell clones that were taken for expansion and analysis.

### APEX proximity labelling

APEX gene expression was induced by supplementing the medium with tetracycline for 24 hr. (50ng/ml for NES/G3BP1; 100ng/ml for FMR1/FXR1). Labelling activity was induced by supplementing Biotin-phenol (BP, 500 μM, Iris Biotech GmbH, LS-3500) for 60 minutes and H2O2 (1 mM, J.T.Baker 7722-84-1) for 1 min. APEX activity extinguish with quenching solution (QS: sodium azide (10mM, Mallinckrodt, 1953-57), sodium ascorbate (10mM, Sigma-Aldrich, A7631) and Trolox (5mM, Sigma-Aldrich, 238813) in PBS. Then, cells were scraped in PBS, centrifuged at 800 × *g* for 10 min at 4°C, pelleted and lysed in ice-cold RIPA lysis buffer supplemented with cOmplete Protease Inhibitor Cocktail (Roche, 4693116001) and PhosSTOP (Roche, 4906837001) and further supplemented with N-Ethylmaleimide (NEM, 2.5mg/ml, Sigma-Aldrich, E3876) in SUMO assays. Lysates centrifuged at 15,000 × *g* for 10 min at 4°C. Protein concentration was quantified with Bio-Rad Protein Assay Dye Reagent (Bio-Rad, 500-0006). Streptavidin-coated magnetic beads (Pierce™ Streptavidin Magnetic Beads, Thermo-Fisher, 88816) were incubated for pulldown experiments (SA-pulldown) with 500µl of the extract at ration of 100µl beads per 500µg of sample with rotation overnight at 4°C.

### Liquid Chromatography and Mass Spectrometry

LC-MS/MS runs were performed on the EASY-nLC1000 UHPLC (Thermo Scientific) coupled to the Q-Exactive Plus or Q-Exactive HF mass spectrometers (Thermo Scientific) (Scheltema et al., 2014). Peptides were separated with a 50 cm EASY-spray PepMap column (Thermo Scientific) using a water-acetonitrile gradient, with a flow rate of 300 nl/min at 40°c. Peptides were loaded to the column with buffer A (0.1% formic acid) and separated using a 105 min linear gradient of 7-28% buffer B (80% acetonitrile, 0.1% formic). The resolutions of the MS and MS/MS spectra were 70,000 and 17,500 for Q-Exactive Plus, respectively. The resolutions of the MS and MS/MS spectra were 60,000 and 30,000 for the Q-Exactive HF, respectively. The m/z range was set to 300-1700 or 380-1800 Th. MS data were acquired in a data-dependent mode, with target values of 3E+06 and 1E+05 or 5E+04 for MS and MS/MS scans, respectively, and a top-10 method.

### Raw proteomic data processing

Raw MS data were processed using MaxQuant version 1.6.2.6(Cox and Mann, 2008). Database search was performed with the Andromeda search engine (Cox et al., 2011) using the human Uniprot database. Forward/decoy approach was used to determine the false discovery rate and filter the data with a threshold of 1% false discovery rate (FDR) for both the peptide-spectrum matches and the protein levels. The label-free quantification (LFQ) algorithm in MaxQuant (Cox et al., 2014) was used to compare between experimental samples, except for the negative controls. Additional settings included carbamidomethyl cysteine as a fixed modification and methionine oxidation and N-terminal acetylation as variable modifications. The “match between runs” option was enabled to transfer identification between separate LC-MS/MS runs based on their accurate mass and retention time after retention time alignment. All raw mass spectrometry data is accessible via ProteomeExchange/ PRIDE repository.

### Proteomics statistical analysis

ProteinGroups output table was imported from MaxQuant to Perseus environment (Tyanova et al., 2016), or R (Team, 2013). Quality control excluded reverse proteins, proteins identified by a single peptide, and contaminants. Non-specific streptavidin-bead binders, were excluded by log2-transformed intensity values. rotein groups required t ≥2 valid values / group. Missing values were replaced by a constant low value. Student’s t test with S0= 0.1 was performed with FDR p-value ≤0.05 for pairs of APEX-On and corresponding APEX-Off samples. Proteins that passed all QC filters were separated for each of the SG-APEX markers and compared to NES samples.. For the stress conditions, LFQ values ≥2 valid values values in at least 1 group. Missing data were replaced using imputation, assuming normal distribution with a downshift of 1.8 standard deviations and a width of 0.4 of the original ratio distribution. Enriched SG proteins were called by student’s t test (SG-APEX vs. NES-APEX) with S0= 0.1 and FDR p-value ≤0.05 and fold-change threshold of Log2 (SG-APEX - NES-APEX) that was based on machine learning that used as training set positive curated SG proteins described in Table S1(FMR1 = 0.96, FXR1 = 0.89, G3BP1 = 0.88). Stress-enriched proteins were tested for association with SG-APEX baits vs. NES-APEX, under basal conditions (without NaAsO2) by student’s t test S0= 0.1 with FDR p-value ≤0.05 and a minimum of 2-fold enrichment (Log2(APEX-SG - NES-APEX) > 1). Temporal SG disassembly study, included proteins that were significantly enriched in FMR1-APEX over NES-APEX values at any of the time points either in GFP or GFP–poly(PR)50 conditions tested as above. Following that, FMR1-APEX values per each group were normalized by the mean of their corresponding NES-APEX values. For analysis of normal SG disassembly, we used ANOVA test with S0=0.1, FDR ≤0.05, standardized by z-score transformation and clustered by Pearson’s correlation coefficients. 2-way ANOVA was used to call proteins that were differentially recruited between GFP and GFP–poly(PR)50 conditions, with FDR ≤0.05. Contrast analysis was used to call proteins enriched in specific time points and between conditions, at FDR ≤0.05. These were standardized by z-score transformations and clustered by Pearson’s correlation coefficients.

### Bioinformatics

The sequences of the proteins that were identified in the MASS-spec experiment were downloaded from uniprot (The UniProt, 2017). Intrinsically disordered regions were predicted using iupred2A (Meszaros et al., 2018) as a stretch of ≥10 amino-acids with IUPred score AA ≥ 0.4, while allowing ≤2 consecutive structured amino-acids (IUPred score of <0.4). Phase separation scores (Pscores) were calculated via (Vernon et al., 2018).

### SG live cell imaging

Cultured cell images were taken by a PCO-Edge sCMOS camera controlled by VisView. installed on a VisiScope Confocal Cell Explorer system (Yokogawa spinning disk scanning unit; CSU-W1 and an inverted Olympus microscope (40× oil objective; excitation wavelengths: GFP - 488 nm; mCherry - 560 nm). SG and cell area were analyzed using surface feature in Imaris software.

### Statistical analysis

Statistics performed with R (Team, 2013). Shapiro-Wilk orLevene tests were used to assess normality of the data. Pairwise comparisons passing normality test were analyzed with atudent’s t-test. Wilcoxon test was used for pairwise comparison of nonparametric data. Multiple group comparisons passing normality test were analysed using ANOVA with post-hoc tests, whereas nonparametric multiple group comparisons were analyzed using the paired-Wilcoxon test when ANOVA assumptions were not met. For analysis of live-imaging disassembly, repeated measures ANOVA test was used with contrast analysis. Statistical P values < 0.05 were considered significant. Data presented as specified in the Figure legends. Data are shown as means ± SEM or SD or graphed using boxplots, as noted in the text.

