## Supplementary materials for "Spatio-temporal Proteomic Analysis of Stress Granule Disassembly Using APEX Reveals Regulation by SUMOylation and Links to ALS Pathogenesis"

#### Supplementary Figures Legends

##### Figure S1 Additional multi-bait calibrations data

(A) Western blot analysis of APEX-G3BP1 and APEX-NES proteins expression, titrated by different concentrations of in growth medium. Tubulin as a loading control. Calibration of APEX-FMR1/FXR1 was done in the same manner.

(B) Western blot analysis of APEX-G3BP1 proteins expression, titrated by different concentrations of in growth medium and compared to endogenous G3BP1 expression (WT-G3BP1 ). Tubulin as a loading control. Calibration of APEX-FMR1/FXR1 was done in the same manner.

(C) Confocal micrographs depicting APEX-NES activity in U2OS cells. Immuno-fluorescence of TUBULIN, neutravidin-Texas-red staining of biotinylated proteins at the proximity of the APEX-NES, that demarcated the soluble cytoplasm and merged channels Lens x63, scale-bar 10µm.

(D, E) Western blot analysis of biotinylated proteins, reflecting of APEX activity after streptavidin pulldown of biotinylated SG proteins. Only endogenously-biotinylated proteins are observed in lanes wherein APEX was not activated (without BP or H2O2).

(F) Confocal micrographs depicting APEX-FMR1 in cells wherein APEX was not activated (without BP). TIA1 immuno-fluorescence, neutravidin-Texas-red staining of biotinylated proteins, and merged xannles. Data demonstrates negligible background when APEX is not active. Lens x63, scale-bar 10µm.

##### Figure S2 Additional analysis pipe-line data of Multi-bait APEX experiment

(A) Analysis pipeline diagram of multi-bait APEX experiment (related to data shown in Figure 1 and 2).

(B) Scatter plot and pearson correlation coefficient values in betwen experimental replicates for any of the four different APEX markers in stress conditions.

(C) Volcano plot of relative protein levels in APEX-On samples relative to APEX-Off samples (x-axis log2 scale) under stress conditions. Y axis depicts the differential expression P values (−log10 scale). Black -proteins not specifically bound to streptavidin beads. Student t-test with correction to multiple hypothesis by FDR. adj. $P < 0.05$.

**Figure S3 Additional analysis pipe-line and data of disassembly APEX experiment**

(A) Analysis pipeline diagram of disassembly APEX experiments (related to data shown in Figure 3 and 4).

(B) Box plot analysis of cell viability under normal, GFP or GFP-PR50 expression. Upper and lower quartiles, and extreme points. ANOVA and Tukey test $P<0.005$.

**Figure S4 Additional data and controls supporting SUMOylation experiments**

(A) Western blot analysis SUMO 1 -conjugated proteins (upper blot) and of biotinylated proteins that serve as loading controls and were developed with streptavidin , after proximity lavbeling of SG with FMR1-APEX activity. Extensive SUMOylation of SG proteins seen as smear at 100-250kD. Representative blot from >3 studies.

(B) Western blot analysis of lysates without or with SUMO ligase inhibitor, 2D08, during stress and recovery, for detection of SUMO 2/3 -conjugated proteins. Representative blot from >3 studies.

(C) Graph bar quantification of SG by live GFP-G3BP1 imaging with increasing concentrations of 2D08 demonstrating that 2D08 treatment does not result in the formation of SG. SG area, normalized to cellular area (y-axis) as a function of time (x-axis). Three experimental repeats for measurement with 4 different areas / well. Representative experiment from > 3 independent live imaging studies.

(D) Western blot analysis of lysates with or without treatment of 2D08 during stress and recovery, for the detection of p-EIF2α and TUBULIN as loading control. The levels of p-EIF2α were comparable across all conditions, demonstrating that 2D08 treatment does not result in phosphorylation of Serine 51 of EIF2α. Representative blot from >3 studies.

**Table S1 data related to multi-APEX bait experiment**

**Table S2 raw data of multi-APEX bait experiment**

**Table S3 data related to SG disassembly experiment**

**Table S4 raw data of multi-APEX bait experiment**

**Table S5 Supplementary materials including antibodies and oligonucleotides**

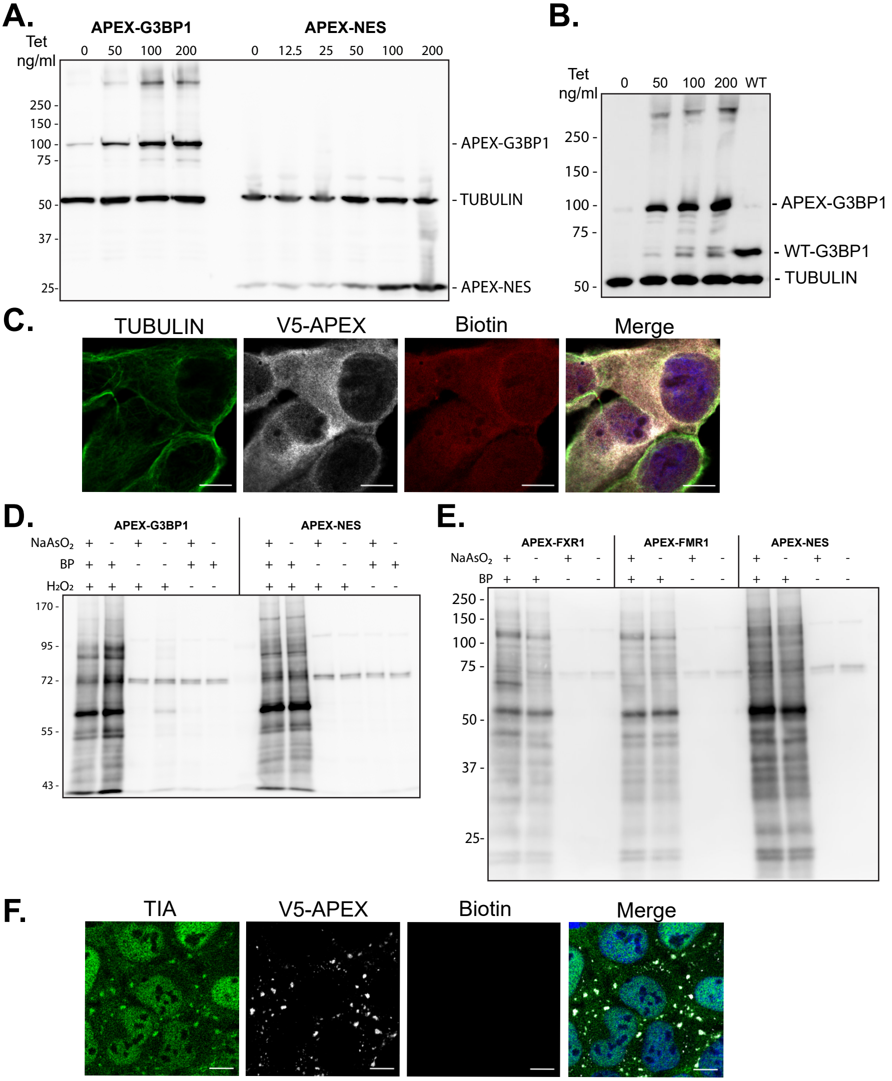


Fig.S1 - Marmor et. al (Hornstein)

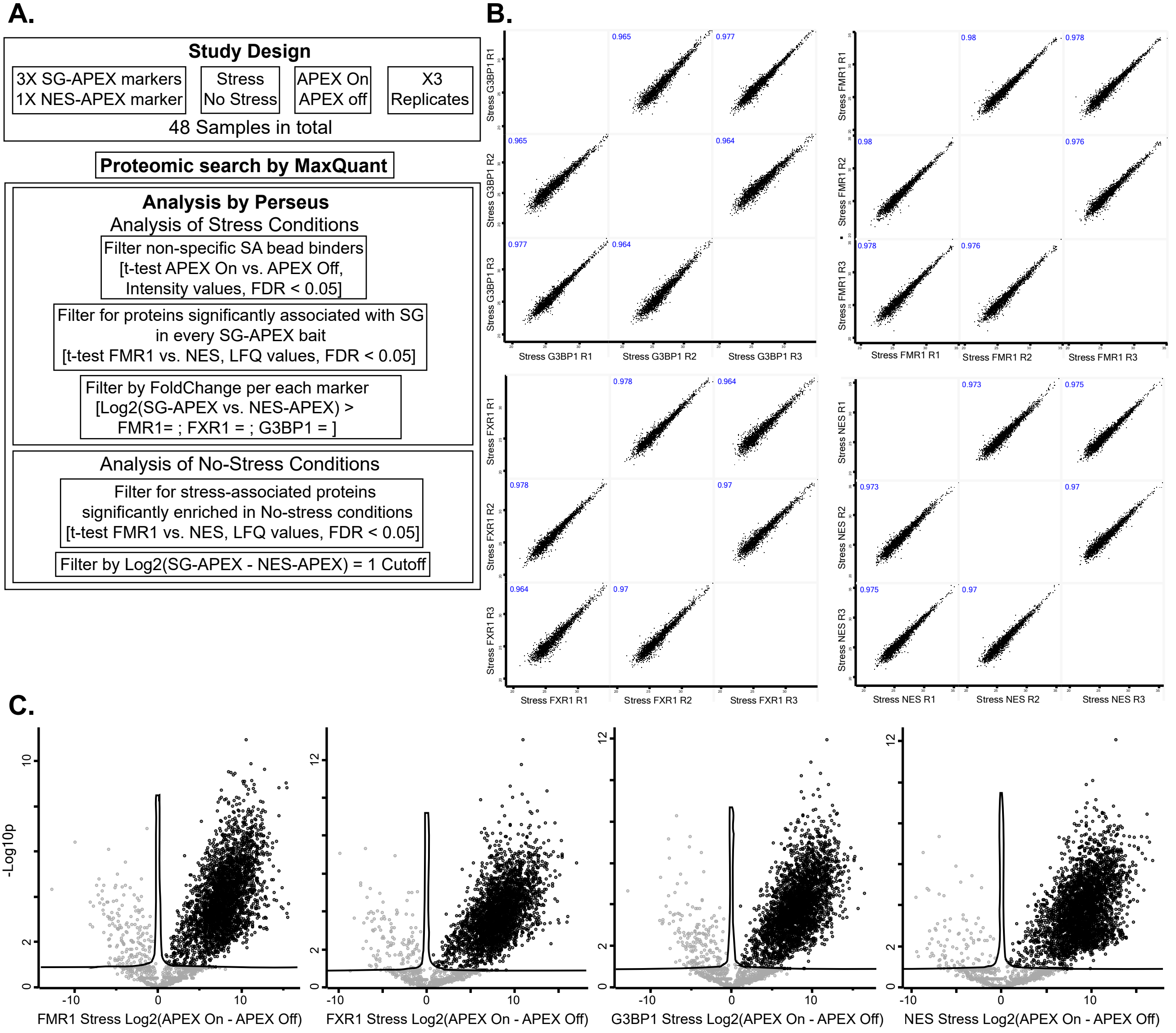


### Fig.S2 - Marmor et. al (Hornstein)

**A.**

#### Study Design

APEX-FMR1; GFP
APEX-NES; GFP
APEX FMR1; GFP-PR50
APEX-NES; GFP-PR50

T0 Stress
T1 60 Min Disassembly
T2 100 Min Disassembly
T3 180 Min Disassembly

APEX On
APEX Off

APEX On - X4 Replicates
APEX Off - X3 Replicates

Total of 112 Samples

#### Proteomic search by MaxQuant

#### Analysis by Perseus

Filter non-specific SA bead-binders
[t-test APEX On vs. APEX Off, Intensity values, FDR < 0.05]

Filter for proteins significantly associated with SG
at any given timepoint or condition
[t-test FMR1 vs. NES, LFQ values, FDR < 0.05, Log2Fold > 1]

Normalize FMR1 LFQ intensities by NES LFQ intensities

##### Analyze Normal SG disassembly Fig. 3D

Analyze GFP samples for proteins
changing in time
[ANOVA, FDR < 0.01, S0=0.01]

##### Analyze differentially recruited proteins GFP-PR50 vs. GFP, Fig. 3F

Analyze GFP and GFP-PR50 samples
for proteins changing between
GFP vs. GFP-PR50, in time
[2-way ANOVA(Time, PR50/GFP),
FDR(PR50/GFP) < 0.05
or FDR(interaction(Time, PR50/GFP)) < 0.05]

Determine internal significance
in each time between GFP and GFP-PR50
[Contrast(GFP/PR50) per each time point,
FDR < 0.05]

**B.**

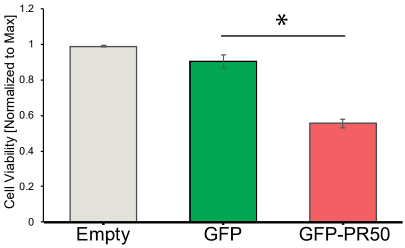


Fig.S3 - Marmor et. al (Hornstein)

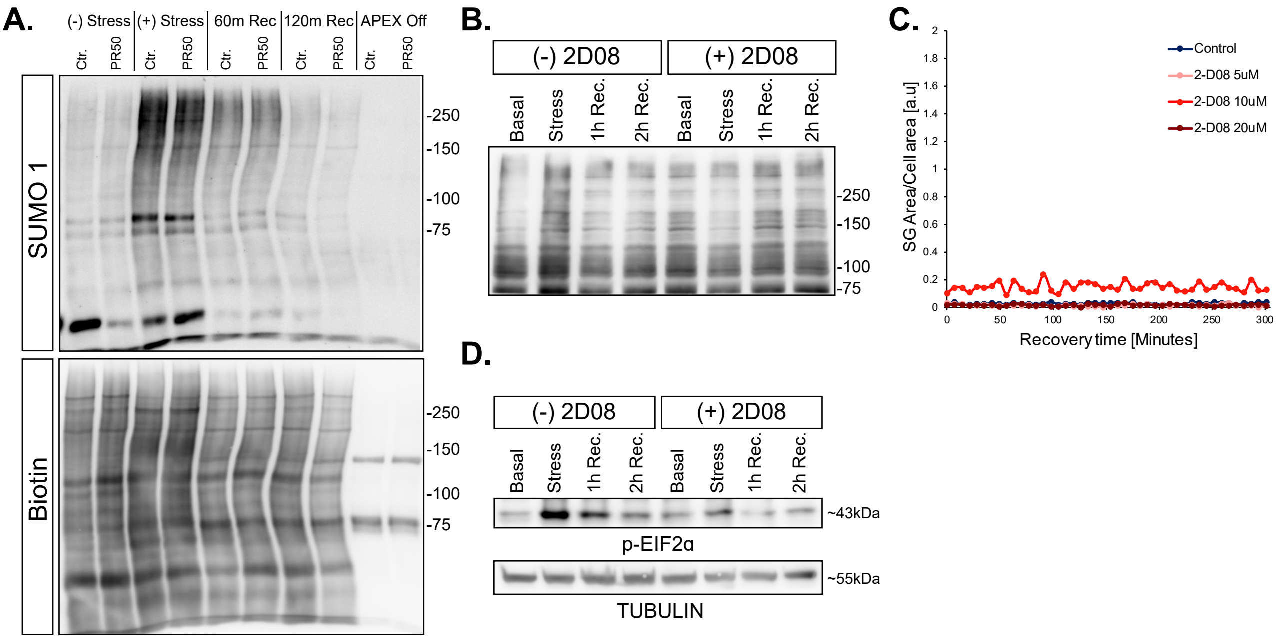


### Fig.S4 - Marmor et. al (Hornstein)
